# Monitoring G-quadruplex formation with DNA carriers and solid-state nanopores

**DOI:** 10.1101/788513

**Authors:** Filip Bošković, Jinbo Zhu, Kaikai Chen, Ulrich F. Keyser

## Abstract

G-quadruplexes (Gq) are guanine-rich DNA structures formed by single-stranded DNA. They are of paramount significance to gene expression regulation, but also drug targets for cancer and human viruses. Current ensemble and single-molecule methods require fluorescent labels, which can affect Gq folding kinetics. Here we introduce, a single-molecule Gq nanopore assay (smGNA) to detect Gqs and kinetics of Gq formation. We use ~5 nm solid-state nanopores to detect various Gq structural variants attached to designed DNA carriers. Gqs can be identified by localizing their positions along designed DNA carriers establishing smGNA as a tool for Gq mapping. In addition, smGNA allows for discrimination of (un-)folded Gq structures, provides insights into single-molecule kinetics of G-quadruplex folding, and probes quadruplex-to-duplex structural transitions. smGNA can elucidate the formation of G-quadruplexes at the single-molecule level without labelling and has potential implications on the study of these structures both in single-stranded DNA and in genomic samples.

---

DNA is the information carrier in the cell and usually forms a right-handed double helix. Beyond the helix, unusual noncanonical structures can naturally occur in a guanine-rich strand of DNA.^1^ These structures known as G-quadruplexes (Gq) are formed from three G-tetrads by forming hydrogen bonds among guanines and stabilized by monovalent ions. More specifically, Gqs tend to form in [G_3_N_X_]_4_ sequences, where four guanines form so-called G-tetrads and three of these tetrads form the full Gq.^2^ For structure stabilization, monovalent or divalent ions are needed, and most of Gqs have the highest specificity for potassium K^+^.^3^ Over recent years, there are numerous studies that show the paramount significance of Gqs for the regulation of gene expression.^4–7^ Hundreds of thousands of these structures were experimentally found in many vital regions of the human genome including but not limited to gene promoters, splicing sites, 3’ and 5’ untranslated regions (UTRs).^8^ Secondly, Gq is shown to be on the basis of many genetic diseases including cancers.^9,10^ Therefore, they are promising drug targets for cancers and effective targets for Ebola and HIV.^11,12^

The two main questions for Gqs are, first, if they are folded, and secondly, how stable they are.^13,14^ A variety of methods, such as circular dichroism (CD), UV-melting analysis, nuclear magnetic resonance (NMR), polymerase stop assay, dimethyl sulfate (DMS) footprinting are able to distinguish folded and unfolded Gqs.^2^ Even though those methods are widely used they usually require a large amount of sample. In parallel to the already discussed ensemble methods, various single-molecule approaches were applied to study Gqs.^15–17^ Widely used to observe Gqs is single-molecule fluorescence resonance energy transfer (smFRET)^18,19^ but similar to many bulk assays smFRET requires fluorescent labelling. A handful of studies pioneered an alternative single-molecule approach based on detection of Gqs with biological nanopores.^20,21^ Similar studies used the same approach to detect if Gqs are folded with silicon nitride nanopores.^22,23^ Here, detection is based on the physical interaction of Gq with the nanopore, again affecting Gq formation. When monitoring competition of Gq DNA and Watson-Crick DNA,^24–27^ fluorescent labeling might affect Gq formation and hence the ratio between Gq and duplex. Understanding Gq formation in the presence of a complementary, competing strand is of great importance for the understanding of Gq formation especially in double-stranded genomic DNA.

In order to create a label-free approach for the analysis of Gqs we combine solid-state nanopore based resistive-pulse sensing and DNA nanotechnology. Glass nanopores are pulled using a P2000 laser puller (Sutter, USA) and assembled in PDMS chips (detailed protocols are presented in section 1 of the Supporting Information). Depending on the experiments we fabricate glass nanopores with diameters ranging from ~5nm to 15 nm. Smaller pores with diameters in the range of 5 nm are used for direct detection of Gq attached to DNA carrier^28^ while larger diameters ~15 nm allow for investigating the quadruplex-duplex competition with streptavidin-labelled strands.^29^ Characteristics of glass nanopores, including their IV curves, used in this study can be found in section 5 of the Supporting Information. The diameter of glass nanopores is double-checked regularly with a scanning electron microscope (SEM) to confirm the resistance values provide accurate size estimates (Figure S-1 of the Supporting Information).

As proof-of-principle for the functionality of smGNA, we designed 190 short oligonucleotides (position named as 1 to 190) that are complementary to linearized single-stranded M13 DNA carrier (for more details see sections 2.1 and 2.2 of the Supporting Information). Oligonucleotides at desired positions can be replaced with short DNA strands containing Gq-sequences at the 3’ end. The three types of Gqs used in this paper are shown in Figure 1a (full sequences of the oligonucleotides are detailed in section 2.3 of the Supporting Information).We choose Gqs which are known to inhibit HIV-1 integrase (T30695)^30^, and to form in human telomeres (hTel)^31^ and minisatellite tandem repeats (26CEB)^32^. We placed a single Gq in the middle of a DNA carrier (Figure 1b). After assembly of DNA carriers following an established protocol^31^ and purification from unbounded oligonucleotides, the sample is diluted in a measurement buffer that always contains a salt concentration of 4 M LiCl, and 0-100 mM KCl, 10 mM Tris-HCl (pH=8.0), 1 mM EDTA and other small molecules as indicated in the respective parts of the manuscript. A diluted sample is incubated for two hours before nanopore measurement unless stated otherwise. The sample is then added to a solid-state nanopore with a diameter of ~5nm. As expected, the presence of one Gq is indicated by a single additional downward peak at or near the designed position. Figure 1c shows one exemplary ionic current trace. The Gq peak current drop (ΔI_peak_) is determined from each event as it is shown in Figure 3c. Custom-written python-based analysis software firstly detects peaks on DNA and after that calculates the relative position of peaks from the closest end of the DNA carrier (more information for data analysis can be found in section 4 of the Supporting Information).

**Figure 1.**
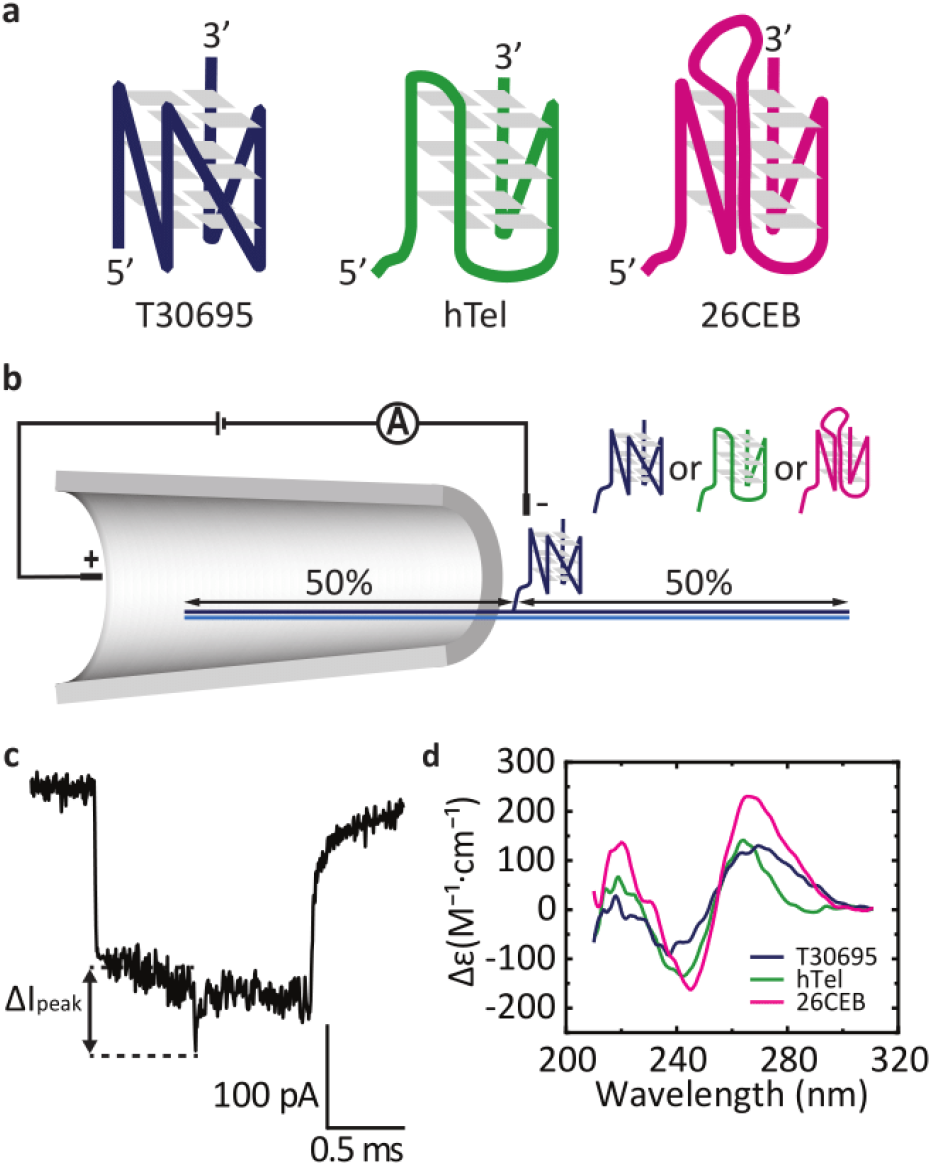
Experimental setup of single-molecule G-quadruplex nanopore assay (smGNA). a) The G-quadruplex structures used in this study are illustrated. b) A DNA carrier with Gq placed in the middle translocates through a nanopore. Other Gqs can be placed in the same position. c) A sample event of the DNA carrier design from b) is shown. The peak in the middle represents T30695 Gq with respective peak current drop (ΔI_peak_). d) Circular dichroism spectra of the three Gq structures in measurement buffer containing 4 M LiCl, 100 mM KCl, 1×TE.

Using bulk measurements, we confirmed the successful formation of all three Gqs using circular dichroism (CD) in the typical measurement buffer (4 M LiCl, 1×TE) with 100 mM KCl (Figure 1d, more information in section 3 of the Supporting Information). Observed CD spectra indicate that Gqs are folded only in presence of 100 mM KCl and in the same conditions with 0 mM KCl Gqs are not folded (Figure S-2), what is in accordance with previously reported data.^26,32,33^ Many groups have investigated the effect of LiCl on Gq folding and find a range of effects both destabilizing^34^ and neutral.^8,17,35^

After confirmation that smGNA can detect one Gq on the DNA carrier, we expand our system. For this purpose, we distribute several Gqs-sequences along the DNA carrier (Figure 2a-c, left panel). Firstly, we designed the DNA carrier with T30695 Gq-sequences ([G_3_T]_4_) at the 3’ end of oligonucleotides in the relative positions at 60, 75, and 87% of the DNA carrier. The nanopores successfully detected our designed construct (Figure 2a, left panel) and relative position of peaks appearing at the designed positions (right panel in Figure 2a). The results for hTel and 26CEB are presented in Figure 2b and 2c, respectively. Again, we observe the Gqs at the expected positions along the carriers. However, the current difference in the current drops among all Gq sequences (Figure 2a-c, middle panel) overlap for all types of Gq. One reason is the variable nanopore diameter. It is important to note that we can clearly detect Gq based on their relative position to one end of a DNA carrier.

**Figure 2.**
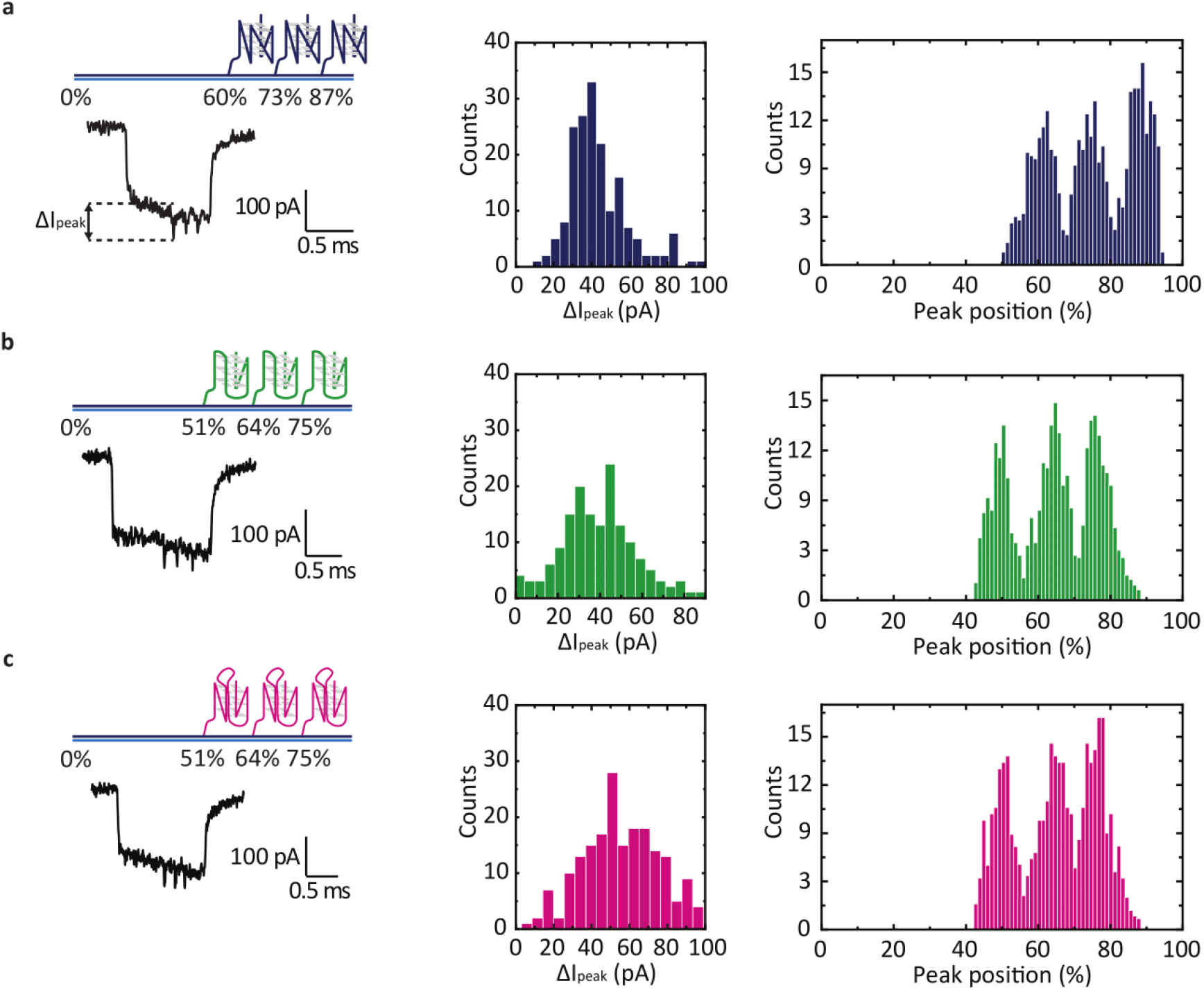
Single-molecule G-quadruplex nanopore assay (smGNA) detects various G-quadruplexes (Gqs) without labelling. Three Gqs are placed at the designed positions as a percentage of the whole DNA carrier (left panel). The current drops □I_peak_ of Gq peaks are shown in the middle. Relative peak positions are shown in the right panel. a) Three peaks of T30695 can be observed with small diameter nanopores (~5 nm). Relative peak positions match the designed positions. The similar design with three Gqs at the different positions is shown for b) human telomere Gq and c) 26CEB Gq.

We further apply smGNA for studying of folding kinetics of T30695 Gq. Firstly, we designed a DNA carrier with six Gqs as shown in Figure 3a. We purified DNA carriers in 10 mM Tris-HCl (pH=8.0), 1 mM EDTA and absence of monovalent and divalent ions. Previous research concluded that in these conditions Gq is not formed.^36^ For the control experiment, we incubated DNA carrier in measurement buffer with 100 mM KCl, to ensure that all Gqs are folded. The nanopore measurement is done in the same measuring buffer containing 100 mM KCl. Obtained data match the expectations, we observe six downward peaks each corresponding to one folded Gq (Figure 3a, Figure S-5 of the Supporting Information).

**Figure 3.**
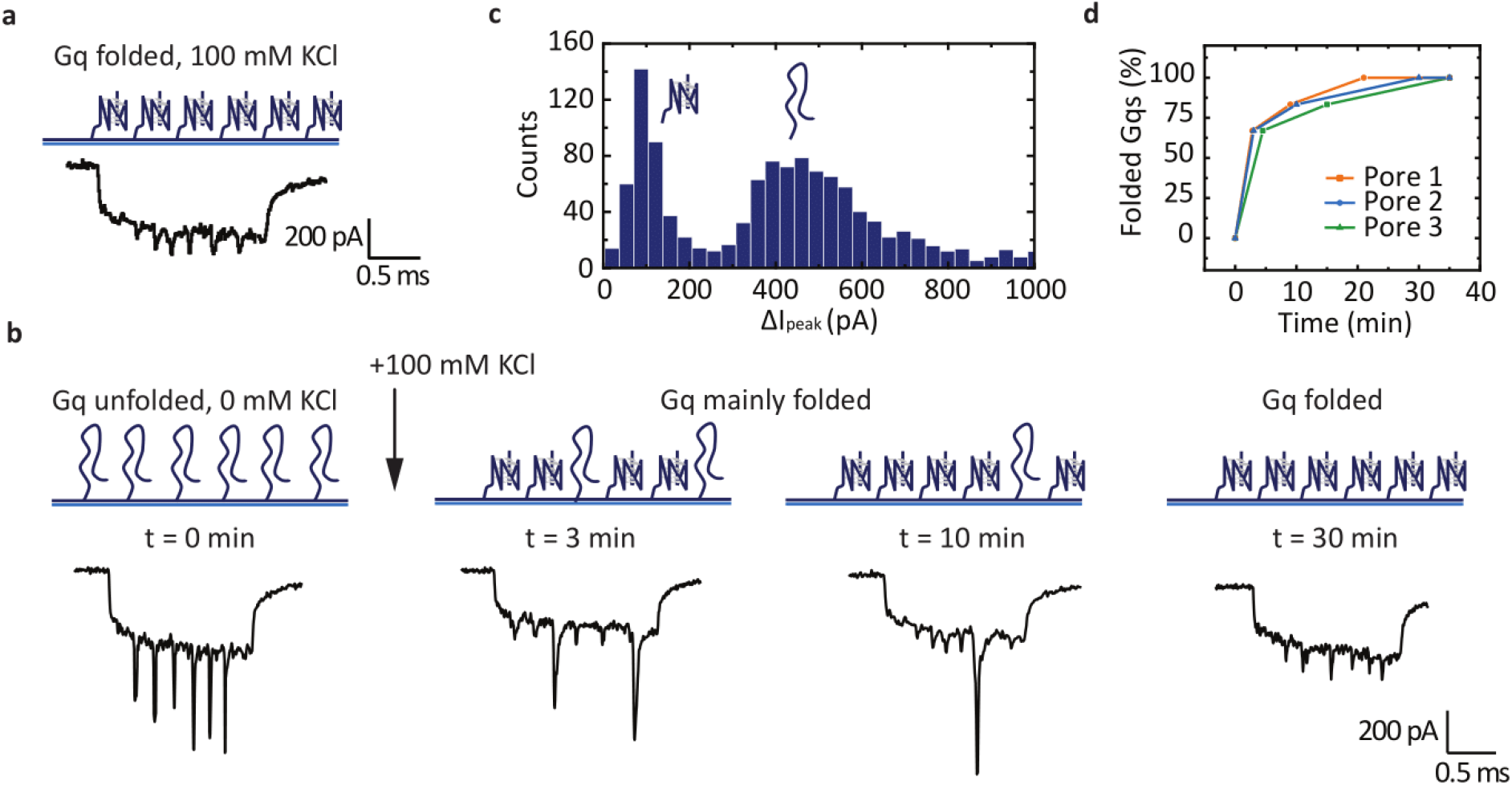
smGNA discriminates unfolded and folded G-quadruplex (Gq). Each peak represents one T30695 Gq attached to DNA carrier. a) Gq pre-folded in 100 mM potassium chloride (KCl) in a nanopore measurement shows only folded Gqs. b) Unfolded Gq in the absence of KCl induces deep peaks at the beginning of the measurement indicate unfolded GQ. Upon addition of unfolded Gqs in measurement buffer with 100 mM KCl, Gqs start to fold. In the first 3 minutes of measurement, the clear majority of Gqs are folded. Nanopores detect unfolded Gq even after 10 minutes. The period of 30 minutes is required for folding of all Gqs. c) We can observe the clear difference between folded and unfolded Gqs from the current drop of peaks (ΔI_peak_). The lower ΔI_peak_ is attributed to folded Gqs and higher current drop are coming from unfolded Gq i.e. single-stranded DNA. d) Single-molecule kinetics of Gq folding for different nanopores is presented.

In order to investigate if smGNA can be used to monitor Gq folding kinetics, we incubated DNA carriers with six bound T30695 Gqs in a measurement buffer with 0 mM KCl. Without potassium we again observe six peaks, but with significantly higher ΔI_peak_ (Figure 3b, t= 0 min). We suspected that these peaks represent unfolded Gqs. To prove that we detect unfolded Gqs, we added 100 mM KCl in solution and continuously monitored DNA carrier translocations over time. It can be observed that after three minutes more than half of Gqs are folded (Figure 3b, t= 3 min). Even after ten minutes, we are still able to detect unfolded Gqs (Figure 3a, t= 10min). Finally, after thirty minutes all Gqs are folded and peaks with higher ΔI_peak_ vanished (Figure 3a, t= 30 min). We confirmed the results by repeating measurements of Gq folding for three different nanopores. For all repeats, we plotted ΔI_peak_ and two distinct populations can be observed (Figure 3c). The population with lower ΔI_peak_ we ascribe to folded Gqs, while the population with higher ΔI_peak_ represents unfolded Gqs. We summarized single-molecule Gq folding kinetics for the three nanopores in Figure 3d. The curves were fitted to one-phase exponential decay function using OriginPro 2018. Observed values for folding rate constant for pores 1, 2, and 3, are 0.39, 0.37, and 0.27 min^−1^, respectively. The differences in the folding rates are within the expected inter-nanopore variability. As mentioned before, deep peaks are caused by unfolded Gq i.e. ssDNA and low peaks are folded Gqs. These experiments represent the first label-free and translocation-based measurement of Gq folding kinetics for solid-state nanopores.

Finally, we expanded our method by investigating quadruplex-duplex binding competition sensing. For this purpose, we used 5’ biotinylated complementary strands of Gq (complementary strand sequences are shown in Table S-2 of the Supporting Information). In the presence of equimolar concentrations of Gq and complementary strand, we assembled three different DNA carriers for the three Gqs (Figure 4a, Figure S-6 of the Supporting Information). Carriers were purified in 10 mM Tris-HCl, pH= 8.0, 0.5 mM MgCl_2_ and after purification stabilizing solution is added as described in section 2.1 of the Supporting Information. For these experiments, we used larger nanopores (~15 nm) as mentioned before, which are easier and faster to fabricate. For larger nanopores, we have to amplify the signal by binding of monovalent streptavidin to the biotinylated complementary strands. With the larger nanopores smGNA detects duplex-biotin-streptavidin (duplex-sb) complex however label-free detection of Gqs is not possible. In this study, we used a monovalent streptavidin with only one biotin active binding site^37^ which was proven to be easily detectable with larger nanopores.^29^ As before, all carriers have three Gqs attached along the DNA carriers (Figure 4a), and hence we can expect a maximum of three peaks which then represent the presence of duplex-sb. The absence of any of designed peaks we attribute to the presence of folded Gq. Sample events for all three DNA carrier designs are presented in Section 6 of the Supporting Information.

**Figure 4.**
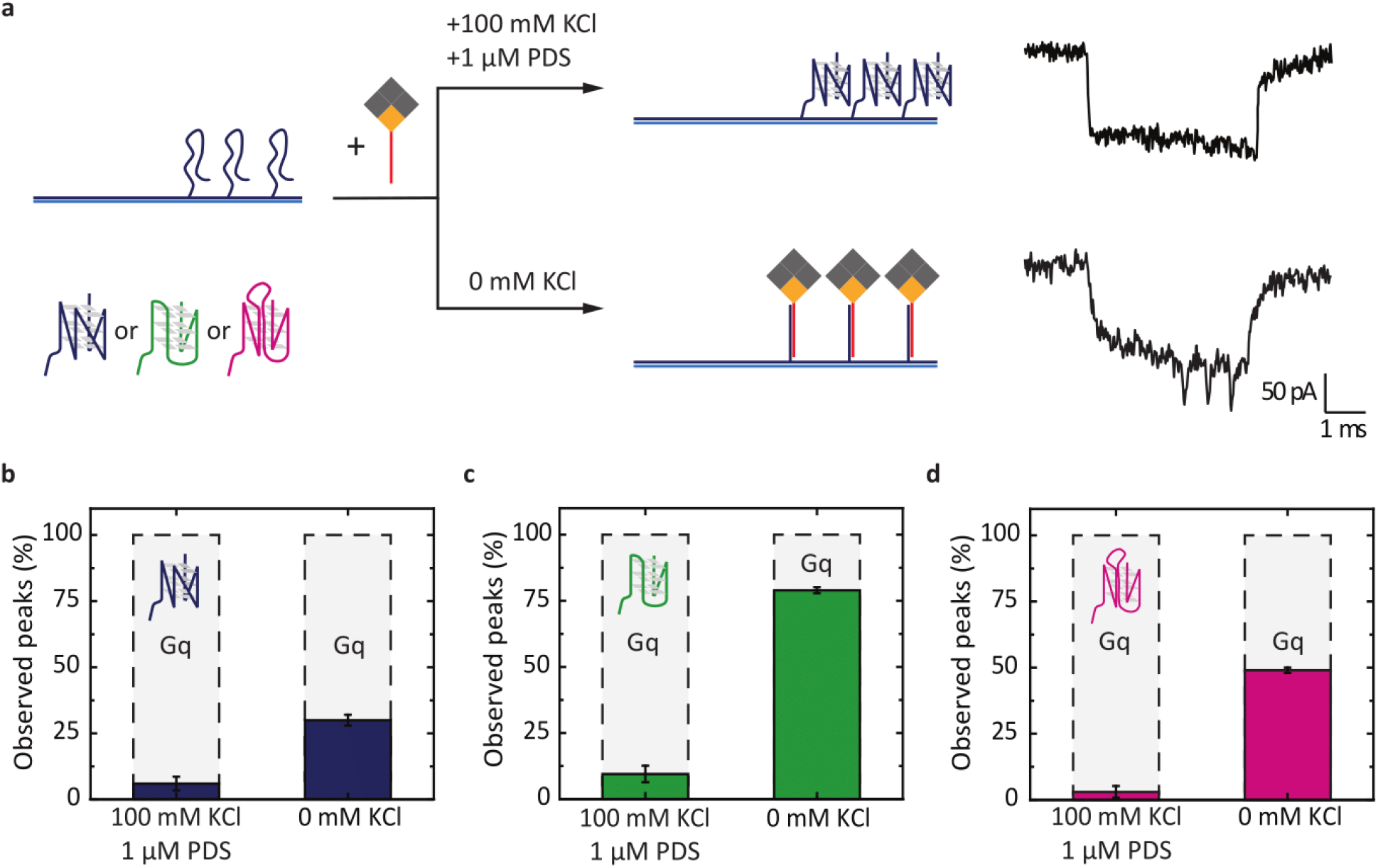
smGNA quantifies quadruplex-duplex competition. a) In the presence of Gq stabilizing conditions including 1 µM high-specific ligand pyridostatin (PDS) and 100 mM KCl majority of Gq are folded (left panel) even in presence of complementary strand in the same concentration. In the absence of stabilizing conditions (0 mM KCl, PDS) duplex forms in different duplex are fomed and peaks are observed indicating duplex formation. The events a) are examples for T30695 in both conditions. b) Histograms of percentages of observed peals for T30695 Gq show that even without stabilizing conditions duplex formation is limited. c) Histogram of observed peaks of hTel duplex formation is significantly promoted without stabilizing conditions and observed peaks are around 79%. d) 26CEB Gq show a similar percentage of folded Gqs as T30695 in stabilizing conditions. For all experiments, Gq and respective complementary strand are mixed in equimolar concentrations. The columns outlined with dashed lines represents folded Gqs. The error bars represent ± standard error mean.

For each of the three Gq structures, two sets of experiments were conducted. In the first set, we favored Gq over duplex formation by assembly of a DNA carrier in Gq-stabilizing conditions i.e. in the presence of 100 mM KCl and 1 µM pyridostatin (PDS). Results indicate that Gq formation is favored in these conditions compared to duplex-sb for all three carriers (Figure 4b-d). We calculated the percentage of observed peaks (duplex-sb) for all three designs and around 90% are in the form of Gq (Figure 4b-d). In the second set of experiments, the same DNA carrier designs were assembled without stabilizing conditions. Example events with three peaks for all three designs are shown in the Supporting Information and the calculated percentage of total observed peaks are presented in Figure 4b-d for direct comparison with Gq formed samples. For all three Gq we see that the number of observed peaks is increasing with removal of KCl and PDS indicating fewer folded Gqs in all cases. Our results are in line with established results as it has been shown that quadruplex-duplex competition can be altered by Gq ligands.^38^ PDS is known as a Gq-stabilizing ligand.^39^ Our results show that, as expected, together PDS and potassium ions prefer Gq over duplex formation.

It is important to note that without potassium ions and PDS, Gqs are still formed. In the case of T30695 Gq, we observed that without stabilizing condition one third of Gq-sequences form duplexes. This is an expected result as for T30695-Gq only in multiple times excess of complementary strand the duplex fully forms.^33^ The hTel Gq without stabilizing conditions indicates increased duplex formation and observed peaks were 79% of all possible peaks. Hence, around 21% of Gqs are folded which is again in accordance with the fluorescent experiments for the same sequence in the presence of 10 mM MgCl_2_.^26^ Finally, 26CEB showed similar results as for T30695, which can be ascribed to almost identical sequence except for the long loop region. Still, the percentage of folded Gqs is smaller compared to T30695. We demonstrated that smGNA can be applied for studying Gq-duplex competition using streptavidin and biotinylated complementary strands as signal amplifiers in larger nanopores. This is a valuable advantage of smGNA for studying Gq formation in dsDNA. smGNA avoids usage of labelling, Gq-binding dyes, and high concentrations compare to other methods applied for detection quadruplex-duplex competition.^26,40,41^

In conclusion, we introduced an innovative single-molecule assay to study G-quadruplex formation in single-stranded DNA and Gq-duplex competition. The smGNA overcomes the necessity of labelling as one of the vital limitations of currently used methods. The smGNA can detect various Gqs and discriminate them based on the designed position along DNA carriers. Secondly, we show that smGNA can discriminate unfolded from folded Gq. Further, we show results for the single-molecule T30695 Gq folding kinetics in the presence of potassium. Finally, smGNA was employed to study Gq-duplex competition. In the presence of stabilizing conditions (100 mM KCl and 1µM PDS) Gq formation is favored over canonical basepairing of a DNA duplex. While without KCl and PDS, equilibrium is shifted closer to duplex formation. Our single-molecule technique will contribute to a better understanding of G-quadruplex formation inside both single-stranded DNA and double-stranded DNA.

## ASSOCIATED CONTENT

### Supporting Information

In detail nanopores fabrication and chip assembly protocol, nanopore statistics, DNA carrier assembly, sequences used in this study, details of CD measurements, sample events and data analysis are presented (PDF).

## Author Contributions

The manuscript was written through the contributions of all authors. All authors have given approval of the final version of the manuscript.

## ACKNOWLEDGMENT

The authors thank Dr Julian Sale and his group, Medical Research Council, Laboratory of Molecular Biology for providing the PDS ligand. The authors thank the Howarth Lab Oxford for the monovalent streptavidin. We a grateful for the help to Catherine Wu for access to CD machines, Department of Chemistry, University of Cambridge. N. Ermann and A. Ohmann are acknowledged for critical reading of the manuscript. F.B., K.C. and U.F.K. acknowledge funding from an ERC Consolidator Grant (Designerpores no. 647144). F.B. acknowledges funding from Benefactors’ Scholarship and scholars’ research expense scheme from St John’s College, Cambridge, United Kingdom, and J.Z. acknowledges funding from an EPSRC grant (EP/M008258/1).

## ABBREVIATIONS

Gq: G-quadruplex
PDS: pyridostatin
duplex-sb: duplex-streptavidin-biotin complex
hTel: human telomere G-quadruplex

