## Supporting Information for "Monitoring G-quadruplex formation with DNA carriers and solid-state nanopores"

### Table of contents

**Section 1. Nanopore chip preparation and measurement conditions**

**1.1. Nanopore fabrication**

**1.2. Nanopore chip assembly**

**1.3. Nanopore diameter estimation**

**1.4. Nanopore measurement**

**1.5. Materials**

**Section 2. DNA carrier preparation**

**2.1. DNA carrier annealing**

**2.2. Oligonucleotides**

**2.3. G-quadruplex sequences**

**Section 3. Circular dichroism recordings**

**Section 4. Data analysis**

**Section 5. Statistics of nanopores**

**Section 6. Sample events**

### Section 1. Nanopore chip preparation and measurement conditions

#### 1.1. Nanopore fabrication

Glass quartz capillaries with filaments with an outer diameter of 0.5 mm and inner diameter 0.2 mm, 7cm in length were purchased from Sutter Instruments, USA. Capillaries were pulled to the desired pore diameter using P2000F model of laser puller (Sutter Instruments, USA). The pulling parameters for 5 ± 1 nm nanopores were[1]:

HEAT=575, FIL=0, VEL=25, DEL=170, PUL=225

For the streptavidin-biotin assay, the inner nanopore diameter was 12 ± 3 nm (mean ± s. d.). This estimation is based on a previously described nanopore fabrication of our method[1,2].

The parameters have to be adjusted to glass quartz capillary batch and are variable between instruments.

#### 1.2. Nanopore chip assembly

Using ceramic blade capillaries were cut to the desired length to connect chip cis and trans chamber. Chip is produced by mixing PDMS and curing agent in the ratio 10:1. Mould was filled with PDMS mixture and left overnight to remove bubbles. The mould with PDMS was solidified by heating on 60°C for 1 hour. The holes in trans chambers were made by biopsy puncher with 1.5 mm diameter and holes between chambers were 1 mm in diameter. Pulled capillaries were placed to connect cis and trans chamber and using plasma etching was attached to a glass slide. The 1 mm holes between cis and trans chambers were filled with 10:1 PDMS mixture and heated for 15 minutes on 100°C to separate possible connections between chambers.

#### 1.3. Nanopore diameter estimation

A glass nanopore diameter is estimated using a scanning electron microscopy (SEM). The inner diameter of nanopores is estimated based on the previous paper[1]. An example of an SEM micrograph is shown below. We used 5 nm and 15 nm nanopores in our experiments. For detection of Gq and kinetics of Gq folding, we used 5 nm nanopores, and for streptavidin-biotin duplex-quadruplex transition, 15 nm nanopores are used. The estimation of nanopore diameter can be made based on the base current. For the 5 nm nanopore at 600mV applied voltage, in 4 M LiCl, 100 mM, 1×TE the base current has to be between 2.94 nA and 4.2 nA. Under the same conditions for 15 nm nanopore, the base current should be between 9 nA and 12 nA.

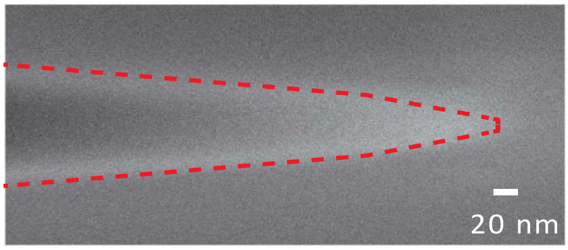


**Figure S-1.** An example SEM micrograph of the glass nanopore used in this study. Scale bar is 20 nm. The tip diameter represents the outer diameter of the nanopore.

#### 1.4. Nanopore measurement

To make glass hydrophilic chip was plasma cleaned for 5 minutes on maximum generator power. Immediately after chambers were filled with working buffer (for instance 4M LiCl, 1×TE). DNA in the concentration of 0.3 nM to 0.5 nM were mixed with the same volume of 8M LiCl, with/without 200 mM KCl, 2×TE, pH=9.44 and then we add 4M LiCl, with/without 100 mM KCl, 1×TE, pH=9.44 (adjusted with 2M LiOH) up to 15 µL. The nanopore measurements were recorded using Axopatch 200B (Molecular Devices, Sunnyvale, CA, USA) and filtered with 50 kHz Bessel filter (Frequency devices, Ottawa, IL, USA). We digitized recordings at a 250 kHz sampling rate with a data card (PCI 6251; National Instruments, Austin, TX, USA).

#### 1.5. Materials

Lithium chloride, lithium hydroxide, potassium chloride, sodium chloride, and 100×TE buffer were purchased from Sigma-Aldrich (1 M Tris-HCl, 0.1 M EDTA, pH 8.0). All buffers were filtered before any experiment using 0.22 µm pore size membrane filter from (Millipore).

### Section 2. DNA carrier preparation

#### 2.1. DNA carrier annealing

The protocol is modified based on the previous paper[2].

Single-stranded DNA carrier – M13 is bought from New English Biolabs - NEB (M13mp18, catalogue number N4040S). As previously described, a DNA carrier with the desired design is prepared as follows:
A 39nt long ssDNA was hybridised to the M13 carrier by mixing
40 µl M13 (stock concentration 250ng/µl),
8 µl 10× NEB cutsmart buffer (Catalog number B7204S),
2 µl of oligonucleotide (100µM, purchased from Integrated DNA technologies - IDT),
28 µl of milli-Q water
Followed by heating to 65°C and cooling to 25°C in a thermocycler over 40 minutes. Then 1 µl of BamHI-high fidelity (R3136T, NEB) and 1 µl of EcoRI-high fidelity (R3101T, NEB) restriction enzymes each at 100000 units/ml, were added to the reaction and incubated for 1 hour at 37°C. The linearized M13 was purified from reaction products using a NucleoSpin gel and PCR clean-up kit (Machery-Nagel), the PCR clean-up protocol was used and the purified M13 is eluted in 2 times 30 µl elution buffer to have high recover of M13.

The next step is annealing of designed oligonucleotides to the carrier. The reaction mixture contained
20.6 µl of mili-Q water,
6.35 µl of cut M13 (126 nM),
4.55 µl of oligonucleotide mixture (527.5 nM of each oligonucleotide),
5.6 µl of 100mM $MgCl_2$,
2.9 µl of 10×TE (100 mM TRIS-HCL (pH=8), 10 mM EDTA)

And heated to 85°C for the 30s, and then on 84.5°C followed by cooling over 1 hour to 25°C. The oligonucleotides are at 3 times excess to the carrier. Nonannealed oligonucleotides were removed using Amicon Ultra 0.5 ml 100kDa filters. One tube of reaction (40 µl) was added to 460 µl of washing buffer (10 mM Tris-HCl (pH = 8.0), 0.5 mM $MgCl_2$) and centrifuged at 6000g for 10 minutes at 4°C. This process is repeated three times. After oligonucleotides leftover removal, around 30 µL of purified DNA nanostructure is collected and 3 µL of stabilizing solution was added (100 mM NaCl, 2 mM $MgCl_2$, 10 mM Tris-HCl, pH=8.0). The concentration of DNA is measured on the Nanodrop UV/Vis spectrophotometer.

*For preparation of the DNA carriers used in direct detection of Gqs (Figure 1-3), purification is done in only filtered 10 mM Tris-HCl solution and stabilizing solution was not added.

To cover M13 carrier with a length of 7228 nt after restriction digestion, 190 oligonucleotides were hybridized (Section 2.2), and desired modified oligonucleotides were in 6 times excess to the carrier. Each oligonucleotide was 38nt long except the last and the first with 46nt in length.

#### 2.2. Oligonucleotides

Below are shown oligonucleotides used to anneal to carrier M13.

**Table S-1.** Oligonucleotide sequences added to anneal a DNA carrier.

| Oligonucleotide number | Sequence (5'-3') | Length (nt) |
|---|---|---|
| 1 | TTTTCGTAATCATGGTCATAGCTGTTTCCTGTGTGAAATTGTTATC | 46 |
| 2 | CGCTCACAATTCCACACAACATACGAGCCGGAAGCATA | 38 |
| 3 | AAGTGTAAAGCCTGGGGTGCCTAATGAGTGAGCTAACT | 38 |
| 4 | CACATTAATTGCGTTGCGCTCACTGCCCGCTTTCCAGT | 38 |
| 5 | CGGGAAACCTGTCGTGCCAGCTGCATTAATGAATCGGC | 38 |
| 6 | CAACGCGCGGGGAGAGGCGGTTTGCGTATTGGGCGCCA | 38 |
| 7 | GGGTGGTTTTTCTTTTCACCAGTGAGACGGGCAACAGC | 38 |
| 8 | TGATTGCCCTTCACCGCCTGGCCCTGAGAGAGTTGCAG | 38 |
| 9 | CAAGCGGTCCACGCTGGTTTGCCCCAGCAGGCGAAAAT | 38 |
| 10 | CCTGTTTGATGGTGGTTCCGAAATCGGCAAAATCCCTT | 38 |
| 11 | ATAAATCAAAAGAATAGCCCGAGATAGGGTTGAGTGTT | 38 |
| 12 | GTTCCAGTTTGGAACAAGAGTCCACTATTAAAGAACGT | 38 |
| 13 | GGACTCCAACGTCAAAGGGCGAAAAACCGTCTATCAGG | 38 |
| 14 | GCGATGGCCCACTACGTGAACCATCACCCAAATCAAGT | 38 |
| 15 | TTTTTGGGGTCGAGGTGCCGTAAAGCACTAAATCGGAA | 38 |
| 16 | CCCTAAAGGGAGCCCCCGATTTAGAGCTTGACGGGGAA | 38 |
| 17 | AGCCGGCGAACGTGGCGAGAAAGGAAGGGAAGAAAGCG | 38 |
| 18 | AAAGGAGCGGGCGCTAGGGCGCTGGCAAGTGTAGCGGT | 38 |
| 19 | CACGCTGCGCGTAACCACCACACCCGCCGCGCTTAATG | 38 |
| 20 | CGCCGCTACAGGGCGCGTACTATGGTTGCTTTGACGAG | 38 |
| 21 | CACGTATAACGTGCTTTCCTCGTTAGAATCAGAGCGGG | 38 |
| 22 | AGCTAAACAGGAGGCCGATTAAAGGGATTTTAGACAGG | 38 |
| 23 | AACGGTACGCCAGAATCCTGAGAAGTGTTTTTATAATC | 38 |
| 24 | AGTGAGGCCACCGAGTAAAAGAGTCTGTCCATCACGCA | 38 |
| 25 | AATTAACCGTTGTAGCAATACTTCTTTGATTAGTAATA | 38 |
| 26 | ACATCACTTGCCTGAGTAGAAGAACTCAAACTATCGGC | 38 |
| 27 | CTTGCTGGTAATATCCAGAACAATATTACCGCCAGCCA | 38 |
| 28 | TTGCAACAGGAAAAACGCTCATGGAAATACCTACATTT | 38 |
| 29 | TGACGCTCAATCGTCTGAAATGGATTATTTACATTGGC | 38 |
| 30 | AGATTCACCAGTCACACGACCAGTAATAAAAGGGACAT | 38 |
| 31 | TCTGGCCAACAGAGATAGAACCCTTCTGACCTGAAAGC | 38 |
| 32 | GTAAGAATACGTGGCACAGACAATATTTTTGAATGGCT | 38 |
| 33 | ATTAGTCTTTAATGCGCGAACTGATAGCCCTAAAACAT | 38 |
| 34 | CGCCATTAAAAATACCGAACGAACCACCAGCAGAAGAT | 38 |
| 35 | AAAACAGAGGTGAGGCGGTCAGTATTAACACCGCCTGC | 38 |
| 36 | AACAGTGCCACGCTGAGAGCCAGCAGCAAATGAAAAAT | 38 |
| 37 | CTAAAGCATCACCTTGCTGAACCTCAAATATCAAACCC | 38 |
| 38 | TCAATCAATATCTGGTCAGTTGGCAAATCAACAGTTGA | 38 |
| 39 | AAGGAATTGAGGAAGGTTATCTAAAATATCTTTAGGAG | 38 |
| 40 | CACTAACAACTAATAGATTAGAGCCGTCAATAGATAAT | 38 |
| 41 | ACATTTGAGGATTTAGAAGTATTAGACTTTACAAACAA | 38 |
| 42 | TTCGACAACTCGTATTAAATCCTTTGCCCGAACGTTAT | 38 |
| 43 | TAATTTTAAAAGTTTGAGTAACATTATCATTTTGCGGA | 38 |
| 44 | ACAAAGAAACCACCAGAAGGAGCGGAATTATCATCATA | 38 |
| 45 | TTCCTGATTATCAGATGATGGCAATTCATCAATATAAT | 38 |
| 46 | CCTGATTGTTTGGATTATACTTCTGAATAATGGAAGGG | 38 |
| 47 | TTAGAACCTACCATATCAAAATTATTTGCACGTAAAAC | 38 |
| 48 | AGAAATAAAGAAATTGCGTAGATTTTCAGGTTTAACGT | 38 |
| 49 | CAGATGAATATACAGTAACAGTACCTTTTACATCGGGA | 38 |
| 50 | GAAACAATAACGGATTCGCCTGATTGCTTTGAATACCA | 38 |
| 51 | AGTTACAAAATCGCGCAGAGGCGAATTATTCATTTCAA | 38 |
| 52 | TTACCTGAGCAAAAGAAGATGATGAAACAAACATCAAG | 38 |
| 53 | AAAACAAAATTAATTACATTTAACAATTTCATTTGAAT | 38 |
| 54 | TACCTTTTTTAATGGAAACAGTACATAAATCAATATAT | 38 |
| 55 | GTGAGTGAATAACCTTGCTTCTGTAAATCGTCGCTATT | 38 |
| 56 | AATTAATTTTCCCTTAGAATCCTTGAAAACATAGCGAT | 38 |
| 57 | AGCTTAGATTAAGACGCTGAGAAGAGTCAATAGTGAAT | 38 |
| 58 | TTATCAAAATCATAGGTCTGAGAGACTACCTTTTTAAC | 38 |
| 59 | CTCCGGCTTAGGTTGGGTTATATAACTATATGTAAATG | 38 |
| 60 | CTGATGCAAATCCAATCGCAAGACAAAGAACGCGAGAA | 38 |
| 61 | AACTTTTTCAAATATATTTTAGTTAATTTCATCTTCTG | 38 |
| 62 | ACCTAAATTTAATGGTTTGAAATACCGACCGTGTGATA | 38 |
| 63 | AATAAGGCGTTAAATAAGAATAAACACCGGAATCATAA | 38 |
| 64 | TTACTAGAAAAAGCCTGTTTAGTATCATATGCGTTATA | 38 |
| 65 | CAAATTCTTACCAGTATAAAGCCAACGCTCAACAGTAG | 38 |
| 66 | GGCTTAATTGAGAATCGCCATATTTAACAACGCCAACA | 38 |
| 67 | TGTAATTTAGGCAGAGGCATTTTCGAGCCAGTAATAAG | 38 |
| 68 | AGAATATAAAGTACCGACAAAAGGTAAAGTAATTCTGT | 38 |
| 69 | CCAGACGACGACAATAAACAACATGTTCAGCTAATGCA | 38 |
| 70 | GAACGCGCCTGTTTATCAACAATAGATAAGTCCTGAAC | 38 |
| 71 | AAGAAAAATAATATCCCATCCTAATTTACGAGCATGTA | 38 |
| 72 | GAAACCAATCAATAATCGGCTGTCTTTCCTTATCATTC | 38 |
| 73 | CAAGAACGGGTATTAAACCAAGTACCGCACTCATCGAG | 38 |
| 74 | AACAAGCAAGCCGTTTTTATTTTCATCGTAGGAATCAT | 38 |
| 75 | TACCGCGCCCAATAGCAAGCAAATCAGATATAGAAGGC | 38 |
| 76 | TTATCCGGTATTCTAAGAACGCGAGGCGTTTTAGCGAA | 38 |
| 77 | CCTCCCGACTTGCGGGAGGTTTTGAAGCCTTAAATCAA | 38 |
| 78 | GATTAGTTGCTATTTTGCACCCAGCTACAATTTTATCC | 38 |
| 79 | TGAATCTTACCAACGCTAACGAGCGTCTTTCCAGAGCC | 38 |
| 80 | TAATTTGCCAGTTACAAAATAAACAGCCATATTATTTA | 38 |
| 81 | TCCCAATCCAAATAAGAAACGATTTTTTGTTTAACGTC | 38 |
| 82 | AAAAATGAAAATAGCAGCCTTTACAGAGAGAATAACAT | 38 |
| 83 | AAAAACAGGGAAGCGCATTAGACGGGAGAATTAACTGA | 38 |
| 84 | ACACCCTGAACAAAGTCAGAGGGTAATTGAGCGCTAAT | 38 |
| 85 | ATCAGAGAGATAACCCACAAGAATTGAGTTAAGCCCAA | 38 |
| 86 | TAATAAGAGCAAGAAACAATGAAATAGCAATAGCTATC | 38 |
| 87 | TTACCGAAGCCCTTTTTAAGAAAAGTAAGCAGATAGCC | 38 |
| 88 | GAACAAAGTTACCAGAAGGAAACCGAGGAAACGCAATA | 38 |
| 89 | ATAACGGAATACCCAAAAGAACTGGCATGATTAAGACT | 38 |
| 90 | CCTTATTACGCAGTATGTTAGCAAACGTAGAAAATACA | 38 |
| 91 | TACATAAAGGTGGCAACATATAAAAGAAACGCAAAGAC | 38 |
| 92 | ACCACGGAATAAGTTTATTTTGTCACAATCAATAGAAA | 38 |
| 93 | ATTCATATGGTTTACCAGCGCCAAAGACAAAAGGGCGA | 38 |
| 94 | CATTCAACCGATTGAGGGAGGGAAGGTAAATATTGACG | 38 |
| 95 | GAAATTATTCATTAAAGGTGAATTATCACCGTCACCGA | 38 |
| 96 | CTTGAGCCATTTGGGAATTAGAGCCAGCAAAATCACCA | 38 |
| 97 | GTAGCACCATTACCATTAGCAAGGCCGGAAACGTCACC | 38 |
| 98 | AATGAAACCATCGATAGCAGCACCGTAATCAGTAGCGA | 38 |
| 99 | CAGAATCAAGTTTGCCTTTAGCGTCAGACTGTAGCGCG | 38 |
| 100 | TTTTCATCGGCATTTTCGGTCATAGCCCCCTTATTAGC | 38 |
| 101 | GTTTGCCATCTTTTCATAATCAAAATCACCGGAACCAG | 38 |
| 102 | AGCCACCACCGGAACCGCCTCCCTCAGAGCCGCCACCC | 38 |
| 103 | TCAGAACCGCCACCCTCAGAGCCACCACCCTCAGAGCC | 38 |
| 104 | GCCACCAGAACCACCACCAGAGCCGCCGCCAGCATTGA | 38 |
| 105 | CAGGAGGTTGAGGCAGGTCAGACGATTGGCCTTGATAT | 38 |
| 106 | TCACAAACAAATAAATCCTCATTAAAGCCAGAATGGAA | 38 |
| 107 | AGCGCAGTCTCTGAATTTACCGTTCCAGTAAGCGTCAT | 38 |
| 108 | ACATGGCTTTTGATGATACAGGAGTGTACTGGTAATAA | 38 |
| 109 | GTTTTAACGGGGTCAGTGCCTTGAGTAACAGTGCCCGT | 38 |
| 110 | ATAAACAGTTAATGCCCCCTGCCTATTTCGGAACCTAT | 38 |
| 111 | TATTCTGAAACATGAAAGTATTAAGAGGCTGAGACTCC | 38 |
| 112 | TCAAGAGAAGGATTAGGATTAGCGGGGTTTTGCTCAGT | 38 |
| 113 | ACCAGGCGGATAAGTGCCGTCGAGAGGGTTGATATAAG | 38 |
| 114 | TATAGCCCGGAATAGGTGTATCACCGTACTCAGGAGGT | 38 |
| 115 | TTAGTACCGCCACCCTCAGAACCGCCACCCTCAGAACC | 38 |
| 116 | GCCACCCTCAGAGCCACCACCCTCATTTTCAGGGATAG | 38 |
| 117 | CAAGCCCAATAGGAACCCATGTACCGTAACACTGAGTT | 38 |
| 118 | TCGTCACCAGTACAAACTACAACGCCTGTAGCATTCCA | 38 |
| 119 | CAGACAGCCCTCATAGTTAGCGTAACGATCTAAAGTTT | 38 |
| 120 | TGTCGTCTTTCCAGACGTTAGTAAATGAATTTTCTGTA | 38 |
| 121 | TGGGATTTTGCTAAACAACTTTCAACAGTTTCAGCGGA | 38 |
| 122 | GTGAGAATAGAAAGGAACAACTAAAGGAATTGCGAATA | 38 |
| 123 | ATAATTTTTTCACGTTGAAAATCTCCAAAAAAAAAGGCT | 38 |
| 124 | CCAAAAGGAGCCTTTAATTGTATCGGTTTATCAGCTTG | 38 |
| 125 | CTTTCGAGGTGAATTTCTTAAACAGCTTGATACCGATA | 38 |
| 126 | GTTGCGCCGACAATGACAACAACCATCGCCCACGCATA | 38 |
| 127 | ACCGATATATTCGGTCGCTGAGGCTTGCAGGGAGTTAA | 38 |
| 128 | AGGCCGCTTTTGCGGGATCGTCACCCTCAGCAGCGAAA | 38 |
| 129 | GACAGCATCGGAACGAGGGTAGCAACGGCTACAGAGGC | 38 |
| 130 | TTTGAGGACTAAAGACTTTTTCATGAGGAAGTTTCCAT | 38 |
| 131 | TAAACGGGTAAAATACGTAATGCCACTACGAAGGCACC | 38 |
| 132 | AACCTAAAACGAAAGAGGCAAAAGAATACACTAAAACA | 38 |
| 133 | CTCATCTTTGACCCCCAGCGATTATACCAAGCGCGAAA | 38 |
| 134 | CAAAGTACAACGGAGATTTGTATCATCGCCTGATAAAT | 38 |
| 135 | TGTGTCGAAATCCGCGACCTGCTCCATGTTACTTAGCC | 38 |
| 136 | GGAACGAGGCGCAGACGGTCAATCATAAGGGAACCGAA | 38 |
| 137 | CTGACCAACTTTGAAAGAGGACAGATGAACGGTGTACA | 38 |
| 138 | GACCAGGCGCATAGGCTGGCTGACCTTCATCAAGAGTA | 38 |
| 139 | ATCTTGACAAGAACCGGATATTCATTACCCAAATCAAC | 38 |
| 140 | GTAACAAAGCTGCTCATTCAGTGAATAAGGCTTGCCCT | 38 |
| 141 | GACGAGAAACACCAGAACGAGTAGTAAATTGGGCTTGA | 38 |
| 142 | GATGGTTTAATTTCAACTTTAATCATTGTGAATTACCT | 38 |
| 143 | TATGCGATTTTAAGAACTGGCTCATTATACCAGTCAGG | 38 |
| 144 | ACGTTGGGAAGAAAAATCTACGTTAATAAAACGAACTA | 38 |
| 145 | ACGGAACAACATTATTACAGGTAGAAAGATTCATCAGT | 38 |
| 146 | TGAGATTTAGGAATACCACATTCAACTAATGCAGATAC | 38 |
| 147 | ATAACGCCAAAAGGAATTACGAGGCATAGTAAGAGCAA | 38 |
| 148 | CACTATCATAACCCTCGTTTACCAGACGACGATAAAAA | 38 |

| | | |
|---|---|---|
| 149 | CCAAAATAGCGAGAGGCTTTTGCAAAAGAAGTTTTGCC | 38 |
| 150 | AGAGGGGGTAATAGTAAAATGTTTAGACTGGATAGCGT | 38 |
| 151 | CCAATACTGCGGAATCGTCATAAATATTCATTGAATCC | 38 |
| 152 | CCCTCAAATGCTTTAAACAGTTCAGAAAACGAGAATGA | 38 |
| 153 | CCATAAATCAAAAATCAGGTCTTTACCCTGACTATTAT | 38 |
| 154 | AGTCAGAAGCAAAGCGGATTGCATCAAAAAGATTAAGA | 38 |
| 155 | GGAAGCCCGAAAGACTTCAAATATCGCGTTTTAATTCG | 38 |
| 156 | AGCTTCAAAGCGAACCAGACCGGAAGCAAACTCCAACA | 38 |
| 157 | GGTCAGGATTAGAGAGTACCTTTAATTGCTCCTTTTGA | 38 |
| 158 | TAAGAGGTCATTTTTGCGGATGGCTTAGAGCTTAATTG | 38 |
| 159 | CTGAATATAATGCTGTAGCTCAACATGTTTTAAATATG | 38 |
| 160 | CAACTAAAGTACGGTGTCTGGAAGTTTCATTCCATATA | 38 |
| 161 | ACAGTTGATTCCCAATTCTGCGAACGAGTAGATTTAGT | 38 |
| 162 | TTGACCATTAGATACATTTCGCAAATGGTCAATAACCT | 38 |
| 163 | GTTTAGCTATATTTTCATTTGGGGCGCGAGCTGAAAAG | 38 |
| 164 | GTGGCATCAATTCTACTAATAGTAGTAGCATTAACATC | 38 |
| 165 | CAATAAATCATACAGGCAAGGCAAAGAATTAGCAAAAT | 38 |
| 166 | TAAGCAATAAAGCCTCAGAGCATAAAGCTAAATCGGTT | 38 |
| 167 | GTACCAAAAACATTATGACCCTGTAATACTTTTGCGGG | 38 |
| 168 | AGAAGCCTTTATTTCAACGCAAGGATAAAAATTTTTAG | 38 |
| 169 | AACCCTCATATATTTTAAATGCAATGCCTGAGTAATGT | 38 |
| 170 | GTAGGTAAAGATTCAAAAGGGTGAGAAAGGCCGGAGAC | 38 |
| 171 | AGTCAAATCACCATCAATATGATATTCAACCGTTCTAG | 38 |
| 172 | CTGATAAATTAATGCCGGAGAGGGTAGCTATTTTTGAG | 38 |
| 173 | AGATCTACAAAGGCTATCAGGTCATTGCCTGAGAGTCT | 38 |
| 174 | GGAGCAAACAAGAGAATCGATGAACGGTAATCGTAAAA | 38 |
| 175 | CTAGCATGTCAATCATATGTACCCCGGTTGATAATCAG | 38 |
| 176 | AAAAGCCCCAAAAACAGGAAGATTGTATAAGCAAATAT | 38 |
| 177 | TTAAATTGTAAACGTTAATATTTTGTTAAAATTCGCAT | 38 |
| 178 | TAAATTTTTGTTAAATCAGCTCATTTTTTAACCAATAG | 38 |
| 179 | GAACGCCATCAAAAATAATTCGCGTCTGGCCTTCCTGT | 38 |
| 180 | AGCCAGCTTTCATCAACATTAAATGTGAGCGAGTAACA | 38 |
| 181 | ACCCGTCGGATTCTCCGTGGGAACAAACGGCGGATTGA | 38 |
| 182 | CCGTAATGGGATAGGTCACGTTGGTGTAGATGGGCGCA | 38 |
| 183 | TCGTAACCGTGCATCTGCCAGTTTGAGGGGACGACGAC | 38 |
| 184 | AGTATCGGCCTCAGGAAGATCGCACTCCAGCCAGCTTT | 38 |
| 185 | CCGGCACCGCTTCTGGTGCCGGAAACCAGGCAAAGCGC | 38 |
| 186 | CATTCGCCATTCAGGCTGCGCAACTGTTGGGAAGGGCG | 38 |
| 187 | ATCGGTGCGGGCCTCTTCGCTATTACGCCAGCTGGCGA | 38 |
| 188 | AAGGGGGATGTGCTGCAAGGCGATTAAGTTGGGTAACG | 38 |
| 189 | CCAGGGTTTTCCCAGTCACGACGTTGTAAAACGACGGC | 38 |
| 190 | CAGTGCCAAGCTTGCATGCCTGCAGGTCGACTCTAGAGGATCTTTT | 46 |

#### 2.3. G-quadruplex (Gq) sequences

The full modified oligonucleotide sequences with attached G-quadruplex (Gq) sequences and sequences of complementary strands with biotin modifications. The Gq sequence itself is bolded at the table. Number/position of oligonucleotide from the table S-1. which is modified is indicated at the table S-2.

**Table S-2.** G-quadruplex sequences and complementary sequences used in this study.

| G-quadruplex full name | Code | Sequence (5'-3') | Modified oligonucleotide | Length (nt) |
|---|---|---|---|---|
| HIV integrase G-quadruplex | T30695 | AATTAACCGTTGTAGCAATACTTCTTTGATTAGTAATA<br>TTT **GGGTGGGTGGGTGGGT** | 25 | 57 |
| | T30695 | GAAACAATAACGGATTCGCCTGATTGCTTTGAATACCA<br>TTT **GGGTGGGTGGGTGGGT** | 50 | 57 |
| | T30695 | TACCGCGCCCAATAGCAAGCAAATCAGATATAGAAGGC<br>TTT **GGGTGGGTGGGTGGGT** | 75 | 57 |
| | T30695 | TTTTCATCGGCATTTTCGGTCATAGCCCCCTTATTAGC<br>TTT **GGGTGGGTGGGTGGGT** | 100 | 57 |
| | T30695 | CTTTCGAGGTGAATTTCTTAAACAGCTTGATACCGATA<br>TTT **GGGTGGGTGGGTGGGT** | 125 | 57 |
| | T30695 | AGAGGGGGTAATAGTAAAATGTTTAGACTGGATAGCGT<br>TTT **GGGTGGGTGGGTGGGT** | 150 | 57 |
| Human telomere G-quadruplex | hTel | GTAGCACCATTACCATTAGCAAGGCCGGAAACGTCACCT<br>TTT **TAGGGTTAGGGTTAGGGTTAGGG** | 97 | 65 |
| | hTel | TGGGATTTTGCTAAACAACTTTCAACAGTTTCAGCGGAT<br>TTT **TAGGGTTAGGGTTAGGGTTAGGG** | 121 | 65 |
| | hTel | GATGGTTTAATTTCAACTTTAATCATTGTGAATTACCTT<br>TTT **TAGGGTTAGGGTTAGGGTTAGGG** | 142 | 65 |
| Human minisatellite 25CEB motif | 26CEB | GTAGCACCATTACCATTAGCAAGGCCGGAAACGTCACCT<br>TTT **AAGGGTGGGTGTAAGTGTGGGTGGGT** | 97 | 68 |
| | 26CEB | TGGGATTTTGCTAAACAACTTTCAACAGTTTCAGCGGAT<br>TTT **AAGGGTGGGTGTAAGTGTGGGTGGGT** | 121 | 68 |
| | 26CEB | GATGGTTTAATTTCAACTTTAATCATTGTGAATTACCTT<br>TTT **AAGGGTGGGTGTAAGTGTGGGTGGGT** | 142 | 68 |

| | | | | |
|---|---|---|---|---|
| T30695 5'-biotin complementary strand | TC | /5Biosg/TTTACCCACCCACCCACCC | / | 19 |
| hTel 5'-biotin complementary strand | HC | /5Biosg/TTTCCCTAACCCTAACCCTAACCCTAA | / | 27 |
| 26CEB 5'-biotin complementary strand | CC | /5Biosg/TTTACCCACCCACACTTACACCCACCCTT | / | 29 |

**Section 3. Circular dichroism recordings**

G-quadruplexes were purchased from Integrated DNA Technologies and dissolved in water to 100 µM (IDT). For the CD measurement oligos were diluted in the respective buffer. Control samples were diluted in either control buffer (4M LiCl, 1×TE) Gq buffer (4M LiCl, 1×TE, 100 mM KCl). With CD measurements we wanted to see if Gqs are folded in control buffer and afterwards if in Gq buffer they are folded. To do that we diluted Gq oligonucleotides to 10 µM in the respective buffer and leave samples overnight at 4°C. CD measurements were recorded on JASCO J – 810 with a temperature-controlled cuvette holder and incubated/measured for 1h at 20°C. A quartz cuvette with a path length of 1 mm was used. CD spectra were obtained as the average of 5 individual measurements in a range between 210 nm and 400 nm, with data interval 0.5 nm, bandwidth 1 nm, scanning speed 50 nm/min and response time of 1 s. CD spectra of Gqs were normalized to the molar ellipticity using the formula $\Delta\varepsilon(M^{-1}\cdot cm^{-1}) = \theta/(32980*c*l)$[3] where θ represents the CD ellipticity in millidegrees (mdeg), c is the Gq concentration in mol/L, and l is the path length in cm. CD spectra of all Gqs used in this study obtained in control buffer are shown in Figure S-2.

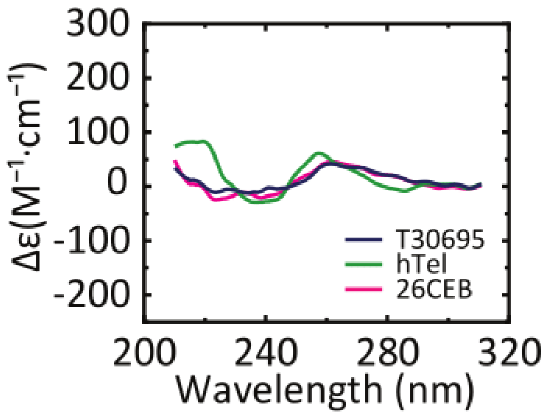


**Figure S-2.** Control CD spectra of Gqs used in this study in 4M LiCl, 1×TE measurement buffer without KCl.

**Section 4. Data analysis**

We used the home-built Python code to find translocations. Since we are collecting single molecule data, short DNA fragments and large aggregates are excluded from further analysis and code distinguish them based on event charge deficit (event surface). Translocation finder isolates events with 100 points before and after an event. Secondly, we set the threshold for peak detection and used Bayesian fitting.[4] Data consist of folded DNA and unfolded DNA. In further analysis, we used only linear-unfolded DNA events.

Positional analysis is done by relative measuring of peak position from the closest end of an event. In this way, we normalize the peak position regardless of the direction in which DNA translocates through a nanopore.

Detail description of nanopore data analysis are presented previously[1–3]. In the case of direct detection of G-quadruplexes, we normalized peak drop to DNA event as shown in Figure 3. This difference is used as **$\Delta I_{peak}$**. All data are plotted using OriginPro 2018 version.

### Section 5. Statistics of nanopores

Here we presented the characteristics of each nanopore measurement we used in this paper including the base current at 600 mV and respective RMS noise.

**Table S-3.** Data for individual nanopores including base current and a root mean square (RMS) noise for the 600 mV voltage used to obtain data.

| Experiment | Nanopore number | Base current at 600 mV (nA) | RMS noise at 600 mV (nA) |
|---|---|---|---|
| Detection of T30695 | 1 | 4.22 | 0.00571 |
| | 2 | 3.71 | 0.00524 |
| | 3 | 3.27 | 0.00559 |
| | 4 | 2.94 | 0.0058 |
| Detection of hTel | 1 | 3.43 | 0.00531 |
| | 2 | 3.42 | 0.00542 |
| | 3 | 3.43 | 0.00555 |
| | 4 | 2.93 | 0.00516 |
| Detection of 26CEB | 5 | 3.2 | 0.00573 |
| | 6 | 3.05 | 0.00564 |
| | 7 | 3.84 | 0.00509 |
| | 8 | 3.6 | 0.00511 |
| | 9 | 3.06 | 0.00508 |
| Kinetics of T30695 Gq folding | 1 | 3.72 | 0.00539 |
| | 2 | 3.12 | 0.00519 |
| | 3 | 3.57 | 0.00545 |
| | 4 | 3.21 | 0.00547 |
| | 5 | 3.82 | 0.00530 |
| | 6 | 3.41 | 0.00593 |
| | 7 | 3.77 | 0.00537 |
| | 8 | 3.31 | 0.00559 |
| | 9 | 3.27 | 0.00556 |
| | 10 | 3.31 | 0.00557 |
| T30965+TC | 1 | 12.59 | 0.00677 |
| | 2 | 10.17 | 0.00676 |
| | 3 | 9.86 | 0.00683 |
| hTel+HC | 1 | 9.28 | 0.00674 |
| | 2 | 9.83 | 0.00660 |
| | 3 | 9.86 | 0.00654 |
| 26CEB+CC | 1 | 9.9 | 0.00659 |
| | 2 | 9.29 | 0.00656 |
| | 3 | 9.13 | 0.00661 |

The IV curves for the nanopores used for data obtaining are shown below. On Figure S-3a. IV curves of nanopores used in direct detection of Gqs and kinetics of Gq folding are plotted. On Figure S-3b. the IV curves used for the quadruplex-duplex competition are plotted.

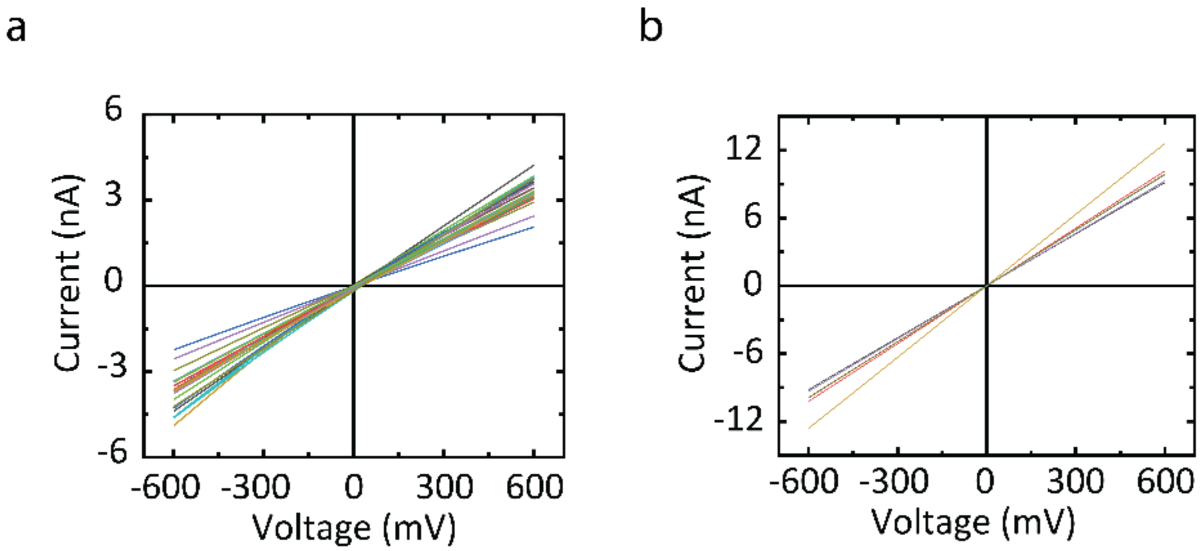


**Figure S-3.** IV curves of nanopores for a) direct detection of Gq and for b) streptavidin-biotin assay.

Example of 2s current trace obtained in this study is shown below. Two events can be observed. The first one unfolded DNA and the second one represents folded DNA.

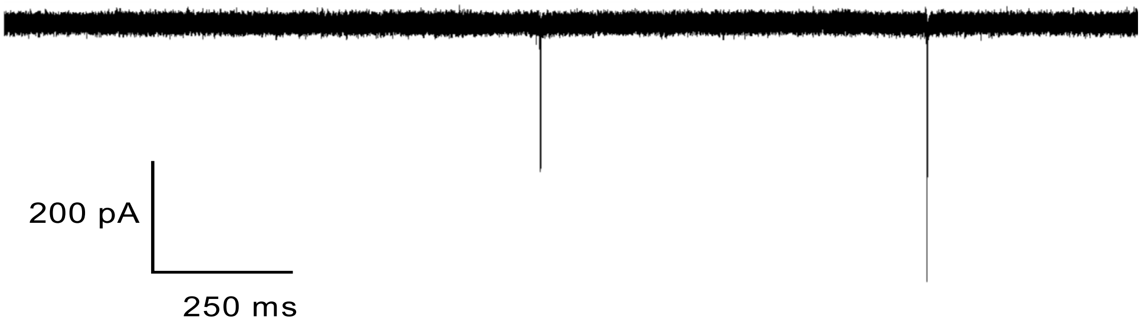


**Figure S-4.** An example of a current trace from the data.

**Section 6. Sample events**

For data presented in Figure 2. and Figure 4. we designed following DNA carriers. Firstly, for T30695 Gq modified oligonucleotides with Gq-sequence were added to 3' end of oligonucleotide positions 25, 50, and 75. Thus, the peaks in positions 25, 50, and 75 corresponds to 13%, 27%, and 40% of event duration, respectively. In Figure 2 peaks positions are shown as 87%, 73%, and 60% (100% minus 13%, 27%, and 40%) for easier graphical understanding of data. Secondly, for hTel and 26CEB Gq modified oligonucleotides with respective Gq-sequence were added to 3' end of oligonucleotide positions 97, 121, and 142. The peaks in positions 97, 121, and 142 corresponds to 51%, 64%, and 75% of event duration. In our data analysis we measure peak position from the closest end of event. In this way we normalize peak position regardless of the DNA carrier direction of entering in nanopore.

The DNA carrier for Gq folding kinetics experiments (Figure 3) had T30695 Gq modified oligonucleotides with Gq-sequence added to 3' end of oligonucleotide positions 25, 50, 75, 100, 125, and 150. In Figure S-4., we show additional sample events for folded T30695 Gqs in 100 mM KCl.

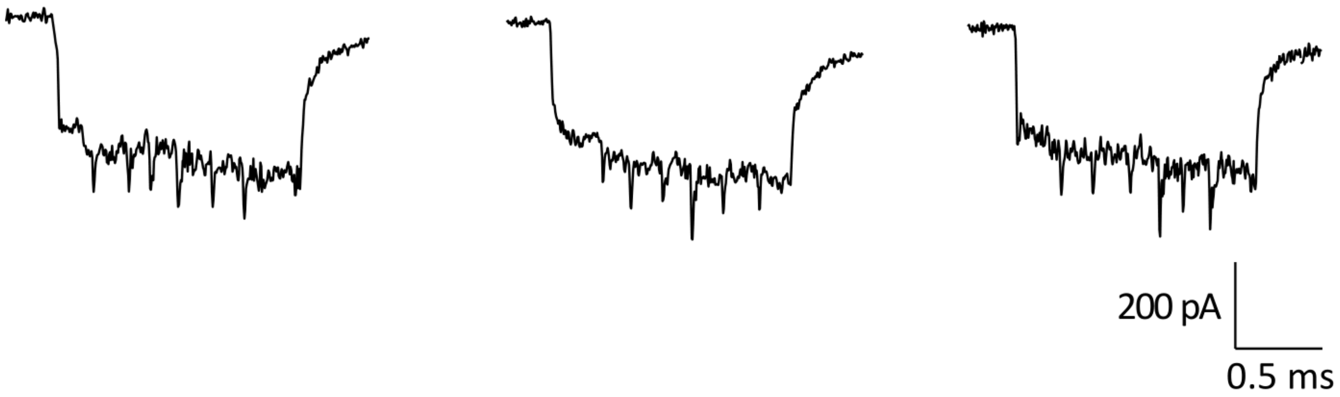


**Figure S-5.** Sample events for folded T30695 Gq in 100 mM KCl are presented. The six peaks corresponding to folded Gq are shown.

For the DNA carrier designs used for quadruplex-duplex competition we exchange one of the 190 oligonucleotides with oligonucleotide with desired Gq-sequence at the 3' end. This was done in step of assembly of DNA carrier as aforementioned. The designed DNA carriers with sample events for both conditions are shown in Figure S-5. For the quadruplex-duplex structural transition data, we have the possibility of a maximum three folded/unfolded Gq so we have either three peaks (all unfolded Gq), two peaks (two unfolded Gq), 1 peak (one unfolded Gq), or no peaks (all folded Gq). Below we show sample events regarding all possible combinations for the quadruplex-duplex competition for T30695 (Figure S-6.), hTel (Figure S-7.) and 26CEB Gq (Figure S-8.). The position is indicated as place where Gq is added to the 3'-sequence of oligonucleotides shown in Table S-1.

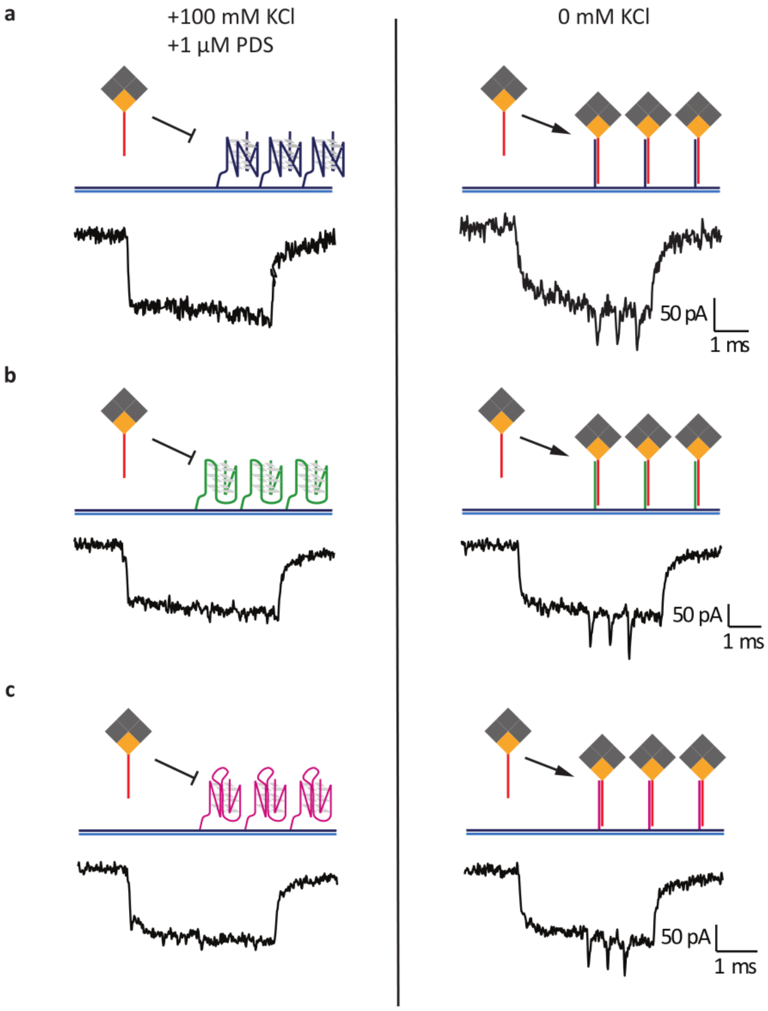


**Figure S-6.** smGNA quantifies quadruplex-duplex competition. In the presence of Gq stabilizing conditions including 1 µM high-specific ligand pyridostatin (PDS) and 100 mM KCl majority of Gq are folded (left panel) even in the presence of complementary strand in the same concentration. In the absence of stabilizing conditions (0 mM KCl) duplex forms in different ratios compare to Gq (right panel). The positions of Gq-forming sequences are identical as in Figure 2. Three Gqs are placed at the specific positions (percentage of the whole DNA carrier). Observed peaks correspond to duplex-biotin-streptavidin (duplex-sb) complex i.e. formed duplex. Here we present the DNA carriers and sample events for a) T30695, b) hTel, and c) 26CEB quadruplex-duplex competition.

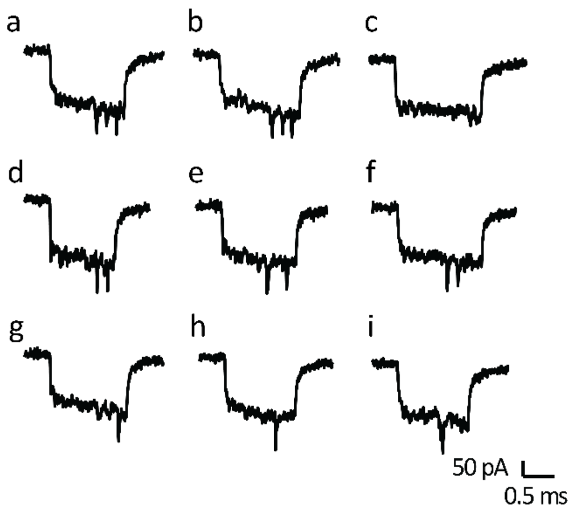


**Figure S-7.** Events as results of the quadruplex-duplex competition for T30695 Gq. Each peak corresponds to duplex with the streptavidin-biotin complementary strand. Three G-quadruplexes are attached to the positions 25, 50 and 75. a), b) all three unfolded, c) all three folded quadruplexes, d) quadruplexes at the positions 25 and 50 unfolded, e) quadruplexes at the positions 25 and 75 unfolded, f) quadruplexes at the positions 50 and 75 unfolded, g) quadruplex at the position 25 unfolded, h) quadruplex at the position 50 unfolded, i) quadruplex at the position 75 unfolded.

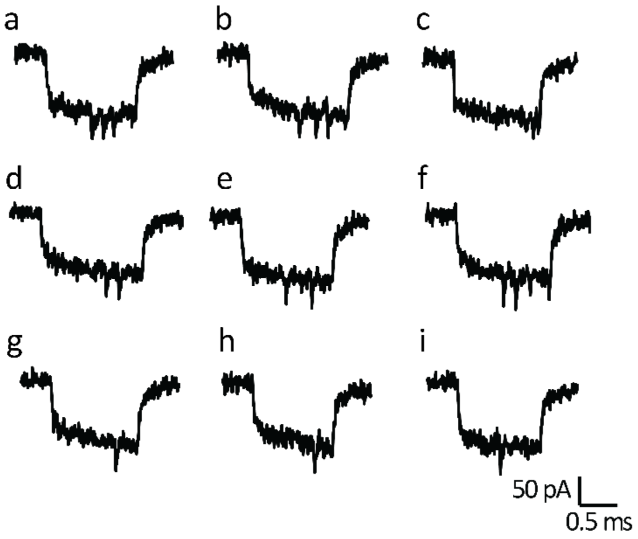


**Figure S-8.** Events as results of the quadruplex-duplex competition for hTel. Each peak corresponds to duplex with the streptavidin-biotin complementary strand. Three G-quadruplexes are attached to the positions 97, 121 and 142. a), b) all three unfolded, c) all three folded quadruplexes, d) quadruplexes at the positions 97 and 121 unfolded, e) quadruplexes at the positions 97 and 142 unfolded, f) quadruplexes at the positions 121 and 142 unfolded, g) quadruplex at the position 142 unfolded, h) quadruplex at the position 121 unfolded, i) quadruplex at the position 97 unfolded.

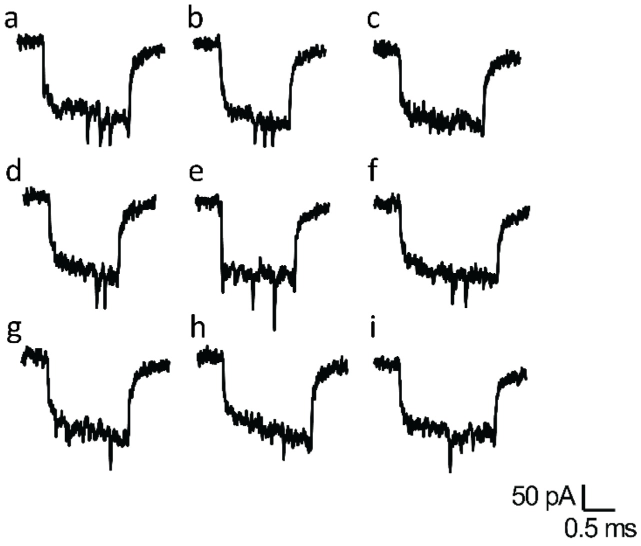


**Figure S-9.** Events as results of the quadruplex-duplex competition for 26CEB. Each peak corresponds to duplex with the streptavidin-biotin complementary strand. Three G-quadruplexes are attached to the positions 97, 121 and 142. a), b) all three unfolded, c) all three folded quadruplexes, d) quadruplexes at the positions 97 and 121 unfolded, e) quadruplexes at the positions 97 and 142 unfolded, f) quadruplexes at the positions 121 and 142 unfolded, g) quadruplex at the position 121 unfolded, h) quadruplex at the position 142 unfolded, i) quadruplex at the position 97 unfolded.
